# Fine mapping chromatin contacts in capture Hi-C data

**DOI:** 10.1101/243642

**Authors:** Christiaan Eijsbouts, Oliver Burren, Paul Newcombe, Chris Wallace

## Abstract

Hi-C and capture Hi-C (CHi-C) are used to map physical contacts between chromatin regions in cell nuclei using high-throughput sequencing. Analysis typically proceeds considering the evidence for contacts between each possible pair of fragments independent from other pairs. This can produce long runs of fragments which appear to all make contact with the same baited fragment of interest. We hypothesised that these long runs could result from a smaller subset of direct contacts and propose a new method, based on a Bayesian sparse variable selection approach, which attempts to fine map these direct contacts.

Our model is conceptually novel, exploiting the spatial pattern of counts in CHi-C data, and prioritises fragments with biological properties that would be expected of true contacts. For bait fragments corresponding to gene promoters, we identify contact fragments with active chromatin and contacts that correspond to edges found in previously defined enhancer-target networks; conversely, for intergenic bait fragments, we identify contact fragments corresponding to promoters for genes expressed in that cell type. We show that long runs of apparently co-contacting fragments can typically be explained using a subset of direct contacts consisting of < 10% of the number in the full run, suggesting that greater resolution can be extracted from existing datasets. Our results appear largely complementary to the those from a per-fragment analytical approach, suggesting that they provide an additional level of interpretation that may be used to increase resolution for mapping direct contacts in CHi-C experiments.

## INTRODUCTION

The three-dimensional structure of the genome influences gene expression at varying levels of scale (Naumova et al., 2013). Multi-megabase compartments of active and inactive chromatin, as well as topologically-associated domains (TADs) spanning hundreds of kilobases, can be readily identified by mapping physical interactions using genome-wide chromatin conformation capture techniques (Hi-C) (Dixon et al., 2012; Van Berkum et al., 2010). However, as Hi-C quantifies interactions between all possible pairs of regions in the genome (e.g. *Hind*III fragments) via massively parallel sequencing, it is inefficient at characterizing individual enhancer-promoter interactions in great depth. To explore such regulatory interactions in detail, the more recently developed Capture Hi-C (CHi-C) method targets sequencing efforts toward interactions between pre-defined regions of interest (“baits”, e.g. *Hind*III fragments overlapping gene promoters) on one end, and all other regions (“prey”) on the other (Jäger et al., 2015; Mifsud et al., 2015a).

CHi-C has enabled identification of contacts made by promoters in primary human cells (Mifsud et al., 2015b; Cairns et al., 2016). The contact maps thus generated show a tendency for multiple contiguous fragments to be linked with the same promoter (Javierre et al., 2016; Burren et al., 2017b), but it is not clear whether enhancers overlapping all these fragments or only a subset of them are directly relevant to the promoter’s regulation. Conversely, the same enhancer region can appear to interact with promoters of multiple genes (Dryden et al., 2014; Martin et al., 2015), while it remains unclear whether this reflects coregulation of these genes. Either phenomenon could also be caused by a lack of resolution in these maps, which are typically constrained by the restriction enzyme used (e.g. *Hind*III produces fragments of median length 4kb). Given that typical enhancers and promoters are considerably shorter than a single *Hind*III fragment (Malin et al., 2013), we hypothesised that collateral contacts may be identified along with the direct enhancer-promoter contacts they neighbour. Such collateral contacts might result from a bait traversing the regions around its primary target via Brownian motion, potentially during the formation of loops (Schwarzer et al., 2016) or from inaccuracies in the cross-linking of proximal regions during the CHi-C procedure (Belmont, 2014; Williamson et al., 2014).

The CHi-C signal around any given bait is represented by counts of read pairs (“counts”) linking that bait fragment to each of its neighbouring prey fragments. The signal exhibits a characteristic exponential decay around the location of the bait (Fig. 1a), thought to reflect Brownian motion rather than biologically interesting interactions. Existing approaches for calling interactions from CHi-C data first fit a regression model to these counts. The model estimates the expected rate of decay by distance from the baited fragment, while accounting for other bait- or prey-specific factors, such as capture efficiency and enrichment bias. Then, the count for each individual bait-prey fragment pair is considered, and those whose count is substantially above that predicted by the regression model are identified (Fig. 1b) (Dryden et al., 2014; Cairns et al., 2016). We noted that CHi-C signals often appeared spatially auto-correlated around interacting prey, in their raw as well as their regression-adjusted form. We sought to use this information to improve resolution of CHi-C contact maps. We hypothesised that joint modelling of neighbouring prey fragments would allow direct contacts to be distinguished from collateral contacts under the assumption that the CHi-C signal peaks at directly contacted preys and gradually decays amongst neighbouring fragments (Fig. 1c).

**Figure 1:**
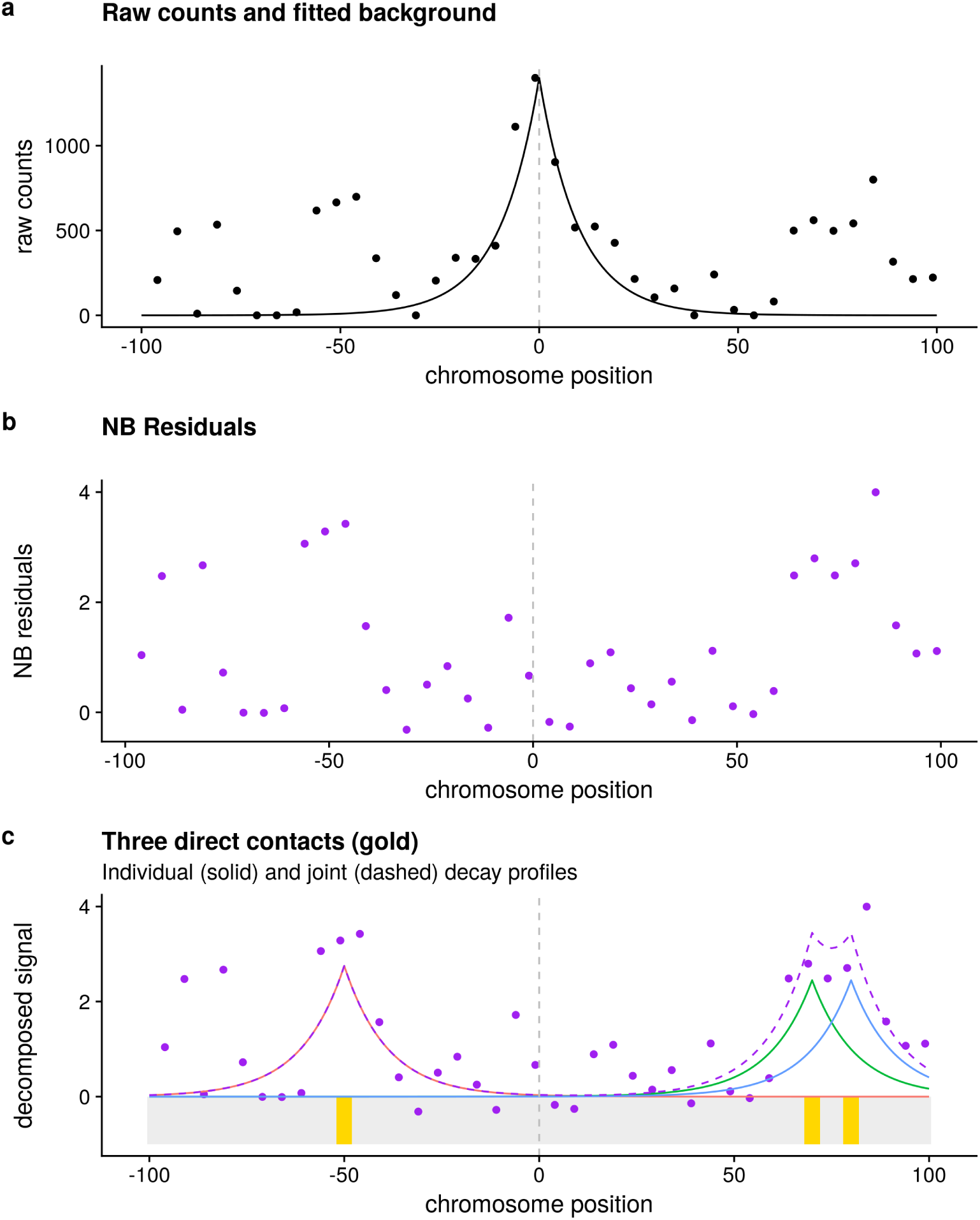
Schematic example of analysis in a single PCHi-C region. a raw counts derived from the PCHi-C experiment and the decay profile around the bait estimated by negative binomial (NB) regression. b residuals after NB regression is used to adjust for expected exponential decay around the bait. c example proposed set of three direct contacts which could be jointly responsible for the spatial distribution of NB residuals, with solid lines indicating the individual decay functions fit in our joint model and the dashed line their predicted joint effect. The position of the bait fragment is indicated by the dotted line and chromosome position is shown in kb relative to the bait.

Here, we propose a statistical model in which, for any given bait, the expected CHi-C signal at each prey is expressed as a sum of contributions from a sparse set of fragments directly contacting that bait. This decomposition model allows us to view the CHi-C signal at each prey in the context of the signals in its local environment. We fit the model through reversible jump Markov Chain Monte Carlo (RJMCMC) to identify primary contacts in published CHi-C data from two cell types, non-activated and activated CD4^+^ T cells (Javierre et al., 2016). Our method prioritises contacts which have greater biological support, suggesting that this model may be useful for increasing precision of contact maps derived from capture Hi-C data.

## RESULTS

### A spatial model for CHi-C data

In common with other approaches (Cairns et al., 2016; Dryden et al., 2014), we first process the read count data from CHi-C using a negative binomial (NB) model to adjust for fragment specific effects as well as the decay in counts around the bait (see Methods). We then define a joint model for the resulting standardised residuals, which we call “NB residuals”. We assume that these NB residuals follow a normal distribution with unit variance (owing to their standardisation) and, in the absence of interactions, zero mean. Where either direct or collateral contacts exist, we propose the NB residuals still follow a normal distribution with unit variance, but with a non-zero mean which we expect is positive. These assumptions are compatible with observed data (see below). Focusing on a single bait, *b*, and its nearest *F_b_* neighbouring prey fragments, let *Y_bp_* be the observed NB residual for prey fragment *p*. Then

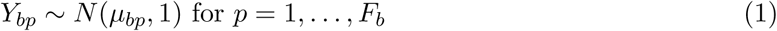

We assume that a direct contact between *b* and a fragment *p* causes μ_*bp*_ to increase by some value *β*_bp_, the magnitude of which reflects the strength of interaction. In the absence of any other contacts we would simply have *μ_bp_* = *β_bp_*. However, we also assume that a direct contact at another fragment, *q*, near to *p*, can affect *μ_bp_*, increasing it by a factor

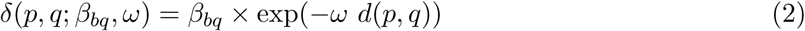

where *β_bq_* captures the strength of the interaction at *q* and *d*(*p, q*) is the absolute linear distance between the midpoints of fragments *p* and *q*, with parameter *ω* (assumed fixed and known) controlling the rate of decay.

The exponential form in (2) was chosen by examining model fits with this and other possible forms of decay functions to a subset of baits. The value of *ω* was fixed at 10^−4.7^, chosen from a range of values tried because it produced the best fit to our data (full details in Supplementary Note, Section 1).

Thus, *μ_bp_* can be expressed as a sum of contributions:

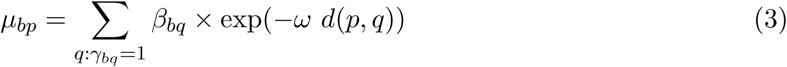

where the sum is taken over all fragments *q* in some neighbourhood of the bait *b*, and *γ_bq_* is a latent indicator variable, taking the value 1 if there is a direct contact between fragments *b* and *q*(i.e. *β_bq_* ≠ 0) and 0 otherwise.

We provide functions to implement this model in the R package Peaky, available from http://github.com/cqgd/pky.

### Inference of bait-prey interactions

We fit the model using an RJMCMC sampler, R2BGLiMS (Newcombe et al., 2014). For each bait *b*, the distribution of sampled coefficients *β*_*b*1_,…, 
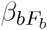
 reflect the posterior distribution of contact strengths between the bait *b* and its neighbouring preys. Given our prior assumption that contacts lead to increased rather than decreased counts, we decided not to use the common marginal posterior probability of inclusion, defined as the proprtion of samples in which *β_bp_* ≠ 0. Instead, we defined an analogous statistic: the marginal posterior probability of a (positive) contact (MPPC) between bait *b* and prey *p*, as the proportion of sampled models in which *β_bp_* > 0 and use this as the primary statistic for inference.

### Application to PCHi-C data from activated and non-activated CD4^+^ T cells

We applied the above model to four parallel data sets generated from CD4^+^ T cells: two from non-activated cells cultured for 4 hours in buffer and two from activated cells cultured for four hours with anti-CD3/CD28 beads, all previously analysed with CHiCAGO (Javierre et al., 2016). Each pair consisted of a *promoter capture set*, with 22,032 bait fragments representing the promoters of 28,007 unique annotated genes (16,116 baits representing 17,731 protein coding genes), and a *validation capture* set, with 945 bait fragments that were preys contacting baits in the promoter dataset according to analysis using the standard CHiCAGO pipeline (Cairns et al., 2016) in CD4^+^ T cells, megakaryocytes or erythroblasts (Javierre et al., 2016). We pre-processed raw counts from each dataset separately to generate NB residuals. QQ plots showed that our assumptions of central normality and a long right tail were met (Supplemental Fig S1). Suggested practice by the authors of CHiCAGO is to declare “significant” interactions when CHiCAGO scores exceed 5 (Cairns et al., 2016). We followed this advice, and focused on baits which had at least one prey fragment with a CHiCAGO score >5 within a window of at least 10 mb around the bait (5 mb either side). We fitted the joint model described by (1) to the NB residuals within these windows for each bait separately in two parallel RJMCMC chains, running additional iterations until the correlation between MPPC from each chain exceeded 0.75. This was achieved for over 94% of baits within 20 million iterations (Table 1, Supplemental Fig S2), and we focus on our inference of these below. The union of samples from both chains was used for inference.

**Table 1:** Number and % of baits for which correlation between MPPC between two parallel runs exceeded 0.75. The total baits is the number for which at least one prey fragment has a CHiCAGO score >5.

| Experiment | Total baits | n. $\rho > 0.75$ | % $\rho > 0.75$ |
| --- | --- | --- | --- |
| Promoter, act | 13684 | 12870 | 94.1 |
| Validation, act | 618 | 583 | 94.3 |
| Promoter, non | 13348 | 12555 | 94.1 |
| Validation, non | 564 | 539 | 95.6 |

### MPPC provides additional information for distinguishing biological plausible contacts

We first compared the MPPC and the CHiCAGO scores for each bait-prey pair. We noted that the CHiCAGO score decayed more rapidly with increasing distance from the bait fragment compared to the MPPC (Supplemental Fig S3), presumably reflecting, in part, the different approaches taken to long-distance contacts. CHiCAGO deliberately down-weights the significance of longer distance (with weights learned based on reproducibility of signals across technical replicates). As our intention is to fine-map longer runs of contacts identified by CHiCAGO, we chose not to apply any down-weighting in order to avoid doubly-penalising them. We also noted that the MPPC and CHiCAGO score were positively correlated (Spearman’s *ρ* > 0.23; Supplemental Fig S4), although a substantial fraction of bait-prey pairs showed high CHiCAGO scores and low MPPC or *vice versa*. We therefore investigated whether one measure alone, or both together, were better at predicting biologically plausible contacts using a variety of measures. We considered that direct contacts from baits in the promoter capture set should be more likely among prey that overlap active chromatin states or that corresponded to published CD4^+^ T cell promoter-enhancer networks (Cao et al., 2017), and in the validation set among prey that contain a gene promoter or the promoter of a more strongly expressed gene. We found that for all of these measures, regression models indicated that either MPPC and CHiCAGO scores together (7 cases) or MPPC alone (1 case) were best able to predict these features (Table 2, Figure 2). This suggested that MPPC and CHiCAGO could be used together to better predict biologically plausible direct contacts than a model which considers each bait-prey pair independently, such as CHiCAGO.

**Table 2:** ΔBIC from the intercept only model for four measures of biological plausibility of contacts. The best fitting model (lowest ΔBIC) is highlighted by *. **a**-**d** are defined in full in the Methods. Briefly, **a** whether the bait-prey pair corresponds to published CD4^+^ T cell promoter-enhancer networks (Cao et al., 2017); **b** whether the prey fragment overlaps active chromatin states defined by Burren et al. (2017a); **c** whether the prey from overlaps a gene promoter; **d** the level of expression of a gene associated with the prey fragment. In all cases, a robust clustered model was used to account for repeated observations at the prey fragment.

| Model | Non-activated | Activated |
| --- | --- | --- |
| **a: Promoter: match to Cao et al.** |  |  |
| MPPC | -387.6 | -249.6 |
| CHiCAGO | -271.2 | -240.9 |
| MPPC + CHiCAGO | * -418.4 | * -304.3 |
| **b: Promoter: link to active chromatin** |  |  |
| MPPC | -785.4 | -402.7 |
| CHiCAGO | -340.4 | -199.0 |
| MPPC + CHiCAGO | * -804.7 | * -417.6 |
| **c: Validation: overlap baited promoter** |  |  |
| MPPC | -429.6 | -285.3 |
| CHiCAGO | -539.3 | -399.7 |
| MPPC + CHiCAGO | * -635.8 | * -457.5 |
| **d: Validation: expression at linked promoter** |  |  |
| MPPC | -1673.2 | * -1297.6 |
| CHiCAGO | -1003.8 | -358.2 |
| MPPC + CHiCAGO | * -1758.9 | -1289.0 |

**Figure 2:**
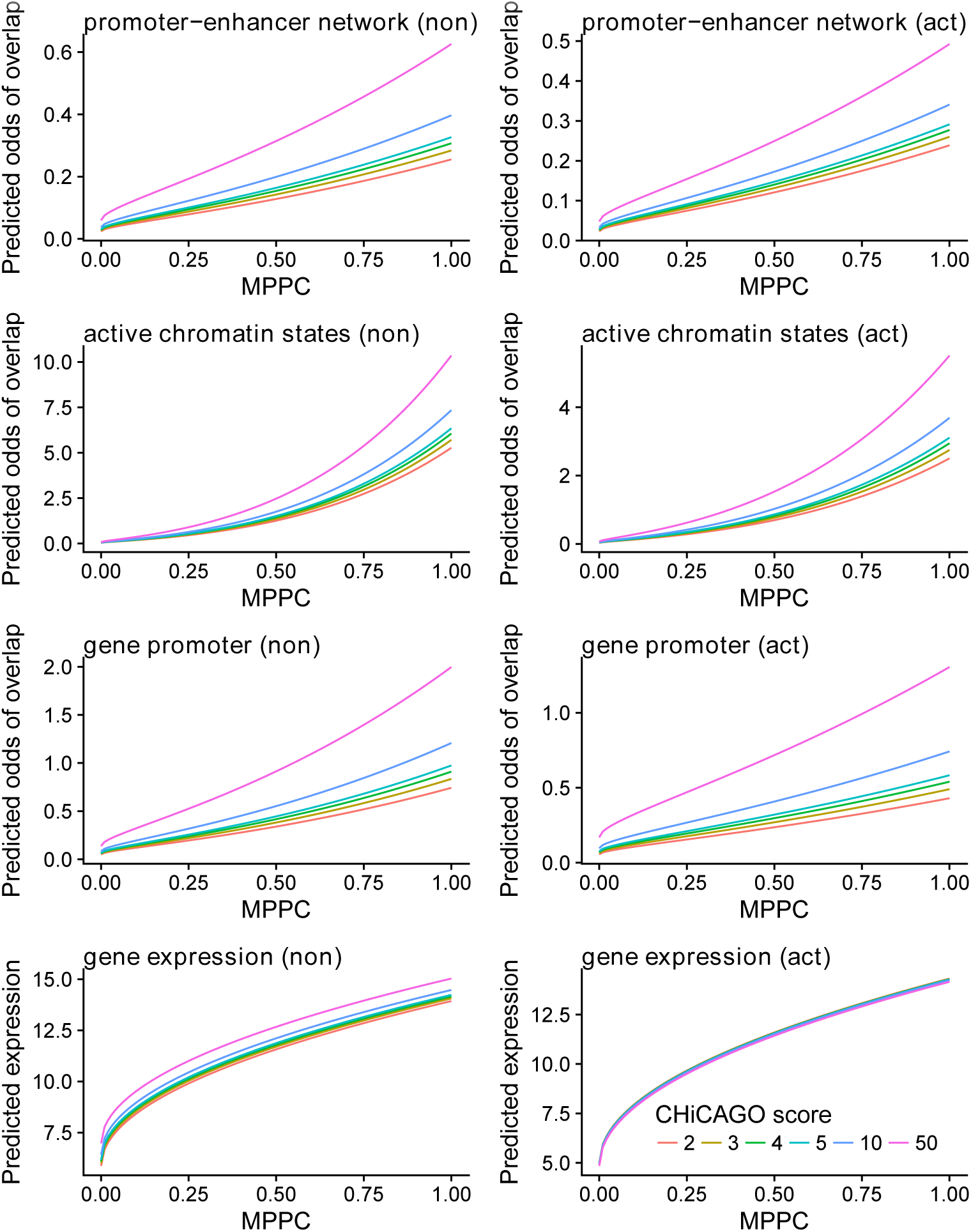
Fitted values from regrssion models of four outcome measures in non-activated (non) and activated (act) CD4^+^ T cells. Row 1: promoter capture, odds that prey fragment overlaps enhancer in published T cell promoter-enhancer network (Cao et al., 2017). Row 2: promoter capture, odds that prey fragment overlaps regions of active chromatin called by CHROMHMM of the same cells (Burren et al., 2017b). Row 3: validation, odds that prey fragment was baited in the promoter capture. Row 4: validation, average gene expression (counts, log_2_ scale) of gene associated with the baited promoter in RNA-seq analysis of the same cell type (Burren et al., 2017b). Predictors are CHiCAGO score (asinh transformed) and MPPC (sqrt transformed).

### MPPC can be used to prioritise direct contacts amongst long runs

The median number of prey fragments per bait identified by a CHiCAGO score > 5 ranged from 7-8, but with a maximum over 200 (Supplemental Fig S5, Supplemental Table S1). In comparison, our model tended to have a slightly greater expected number of contacts per bait when the CHiCAGO count was low (many of these related to fragments with CHiCAGO scores ∈ [3, 5)), but many fewer when the CHiCAGO count was high (Supplemental Fig S5). To enable discussion of longer runs of fragments with high CHiCAGO scores, we define a “stretch” of length *n* to be a series of *n* adjacent fragments with CHiCAGO scores > 5.In the longest stretches of length 50 or more, our model estimated the expected number of direct contacts to be ~ 7% of the number of prey (Supplemental Fig S6). The posterior was spread over a larger number of fragments than the number of expected contacts, but it was not uniform, and we found that the majority of the posterior was often concentrated within a minority of the fragments: for example, within stretches of length 50 or more, a median of 76% of the corresponding posterior space of interaction combinations could be captured by just the top 30% of the fragments ranked by MPPC or 90% of the posterior by the top 50% of the fragments (Fig 3).

**Figure 3:**
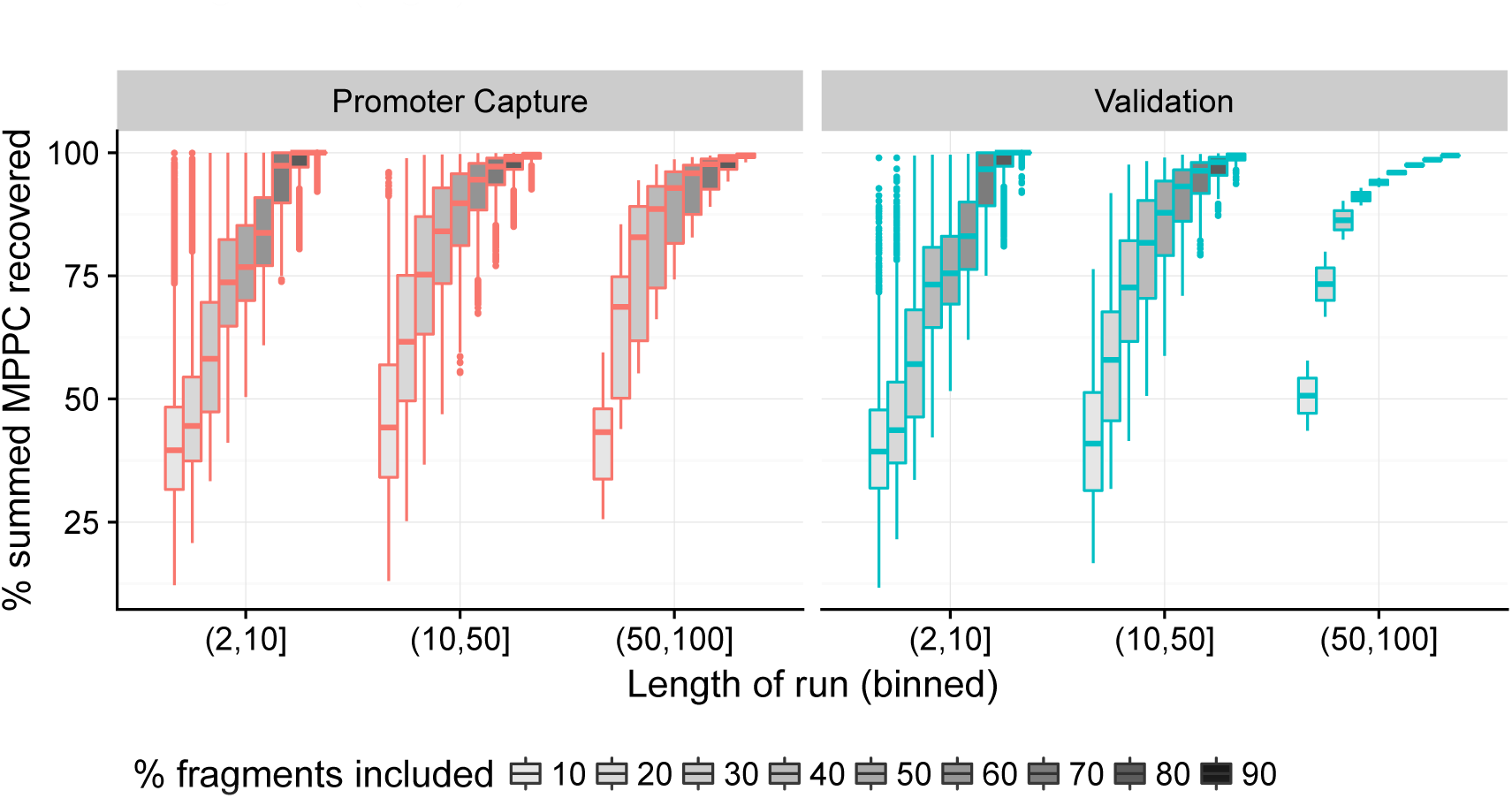
Runs of fragments with CHiCAGO scores > 5 were binned by length (x-axis), and fragments within runs ordered by decreasing MPPC. Boxplots show the proportion of total posterior support captured by including increasingly larger subsets of the ordered fragments. For longer runs (> 10 fragments) the majority (> 50%) of the posterior, as quantified by the summed MPPC across the run, is generally be captured by a minority (< 10%)of the fragments.

This suggested that long stretches could result from direct contacts at a small subset, and that our joint model could distinguish these, ranking some as more probable direct contacts than others. As the true sets of direct contacts are unknown, we again used external data to assess whether this prioritisation corresponded to fragments with more biologically supportable characteristics. We found that, within these stretches, the MPPC remained significant predictors of whether fragments corresponded to biologically plausible features across all run lengths (Supplemental Table S2).

Finally, we illustrate these results by considering example baits with long runs of CHiCAGO-significant prey fragments. First, Fig. 4 shows the analysis in the region of the *ETS1* gene promoter. Here, runs totalling 218 prey fragments have CHiCAGO scores >5 which our joint analysis suggests may be explained by a subset of 7 fragments. Our best estimates of where these fragments lie, highlighted by the peaks in MPPC, clearly correspond to peaks in H3K27ac. Similarly, a bait on chromosome 2 which is annotated with both an antisense gene AC009505.2 and an alternative promoter of *NCK2* is linked to long runs totalling 198 CHiCAGO-significant prey fragments which our model suggests can be explained by subsets of 7 fragments (Fig. 5). Visually, the stretch of fragments surrounding the promoter are split by MPPC into 5 groups, each of which correspond to regions with active chromatin marks.

**Figure 4:**
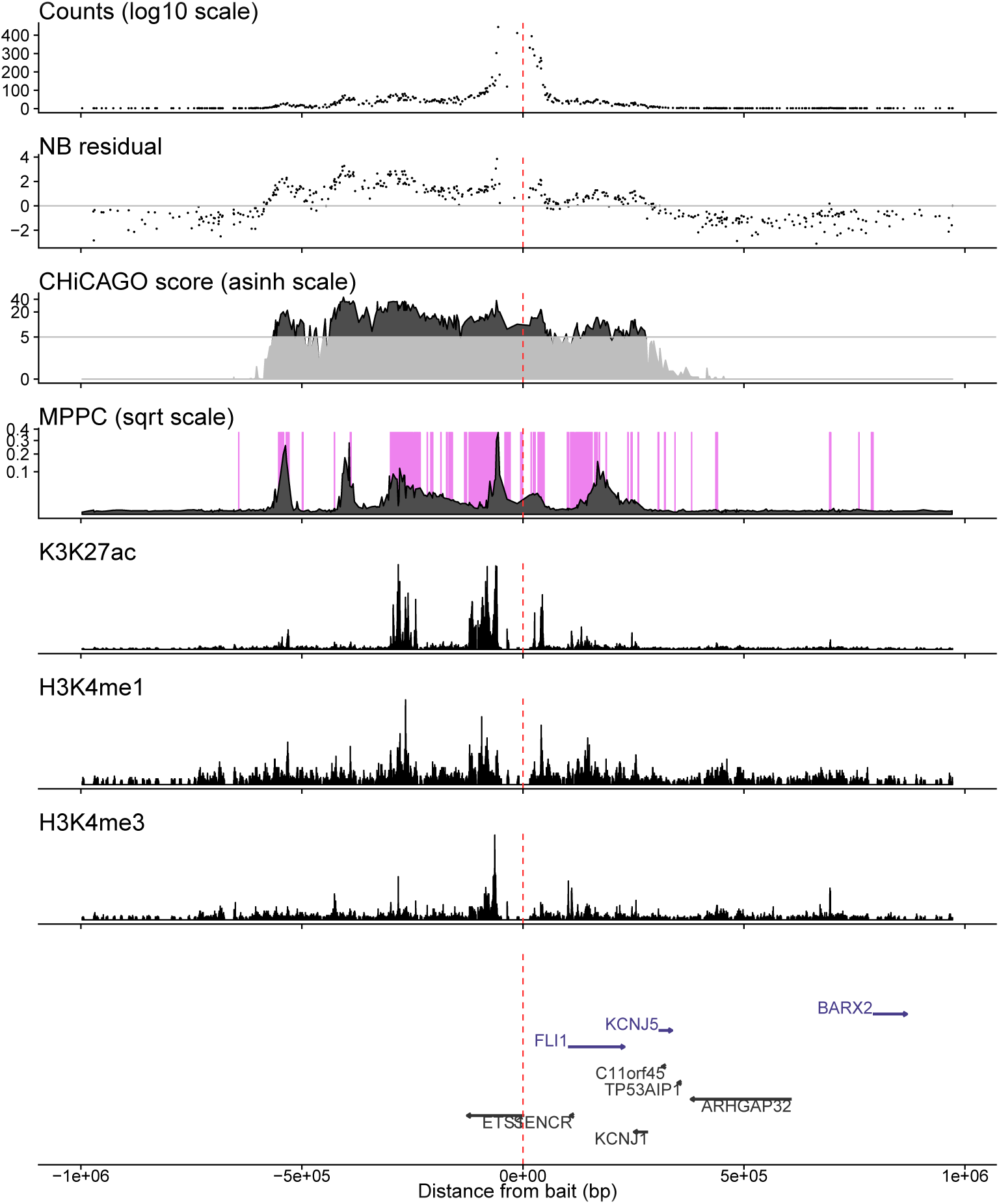
Illustrative example of analysis in the region of the *ETS*1 promoter in activated CD4^+^ T cells. The top three panels show raw counts, adjusted residuals, CHiCAGO scores, and MPPC. The MPPC is overlaid on shading highlighting regions of active chromatin derived by aggregating states from CHROMHMM analysis ChIP-seq data from the same cell type, as previously published(Burren et al., 2017b). The next three panels show three examples of this CHiP-seq data: H3K27ac, H3K4me1, H3K4me3. The final panel shows gene locations. The red vertical line shows the location of the bait fragment. Note that the long run of fragments around and to the left of the promoter with high CHiCAGO scores are split by MPPC into five groups, all of which correspond to regions with active chromatin marks.

**Figure 5:**
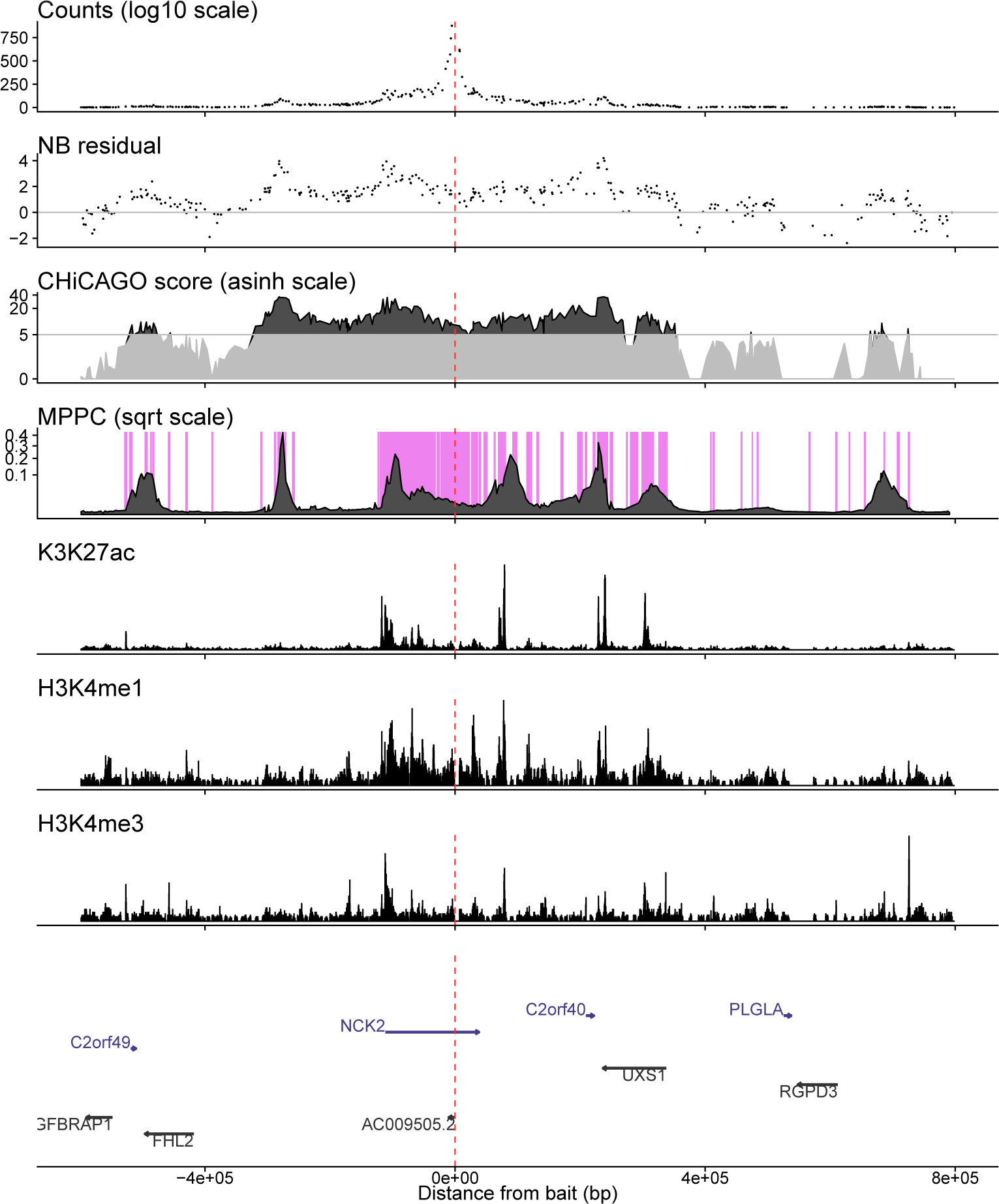
Illustrative example of analysis in the region of the a bait for the antisense gene AC009505.2 and an alternative promoter of *NCK2* in non-activated CD4^+^ T cells. The top three panels show raw counts, adjusted residuals, CHiCAGO scores, and MPPC. The MPPC is overlaid on shading highlighting regions of active chromatin derived by aggregating states from CHROMHMM analysis ChIP-seq data from the same cell type, as previously published (Burren et al., 2017b). The next three panels show three examples of this CHiP-seq data: H3K27ac, H3K4me1, H3K4me3. The final panel shows gene locations. The red vertical line shows the location of the bait fragment. Note that the long run of fragments to the left of the promoter with high CHiCAGO scores are split by MPPC into two groups, which correspond to regions with active chromatin marks.

## DISCUSSION

Our results support our hypothesis that long runs of prey fragments with high counts in PCHi-C data can result from a smaller number of direct contacts together with collateral signal at their neighbours. This suggests that efforts to jointly model the pattern of counts across multiple fragments have potential to distinguish those direct contacts. Joint modelling to improve resolution is already used to fine map genetic causal variants in genome-wide association studies (GWAS). It is accepted in GWAS that the p value corresponding to a test of association between a single genetic variant and some phenotype should be interpreted in the context of the p values of its neighbours, either by highlighting the variant in the region with the smallest p value, or by fitting a variable selection model to find a sparse subset of variants which could explain the association signals across the region. The primary difference between our PCHi-C model and the class of GWAS fine mapping models that also fit the association statistics directly (e.g. PAINTOR, Kichaev et al. (2014)) is that the decay of GWAS association signals across genetic variants has been established to relate to the linkage disequilibrium or correlation between those variants within the population, while our model assumes an exponential decay specified by a single parameter *ω*. We chose *ω* by considering a range of values and choosing that which produced residuals without obvious autocorrelation. This meant we could parallelise our analysis, considering each bait independently, but different values of *ω* would produce different results. Future work will explore whether it is computationally feasible to specify *ω* within a hierarchical framework that considers multiple baits simultaneously. In addition, we intend to investigate whether these ideas – using information from sets of proximal locations in a joint model to make inference about each individual location – could be adapted to other techniques used to call DNA contacts such as ChIA-PET and Hi-ChIP, although different decay functions might be required.

In addition to jointly modelling the signal across multiple fragments, our proposed model contrasts to previous efforts to analyse CHi-C data by producing a Bayesian measure of confidence in the location of a direct contact - the MPPC. Both the MPPC and the CHiCAGO score decay with distance from bait, emphasising that short range contacts predominate, at least within the set of contacts detectable through PCHi-C. There is, though, a notable difference between the rates of decay (Supplemental Fig S3). This reflects the deliberate choice of the CHiCAGO authors to weight p values such that more distant interactions were less likely to be called significant. We chose not to adopt any distant-dependent prior as our intention was to fine map contacts already called by a method that incorporates this distance penalty, such as CHiCAGO, and we did not wish to doubly penalise distant contacts. However, it is possible that adopting such a prior would lead to improved inference were our joint model to be applied alone. We argue that our joint analysis of neighbouring prey fragments adds a further useful dimension to the analysis of capture Hi-C data, with a CHiCAGO score reflecting the (distance from bait-adjusted) evidence for there being any contacts in a neighbourhood, and the MPPC reflecting the expected number and the uncertainty in the precise location of direct contacts. Other advantages of adopting a Bayesian framework include the ability to extend the model to include not just bait-prey distance, but other prior information on the likelihood of direct contacts. This would enable, for example, information from previous experiments in related cell types to inform future analyses.

We have proposed a new model for calling direct contacts from CHi-C data that, in contrast to existing fragment-by-fragment analysis methods, exploits information from each prey’s neighbouring fragments. Our joint model identifies prey fragments with biological characteristics that would be expected at sites of direct contact, such as an active chromatin state when they contact promoters. We have shown this information is largely complementary to that produced by the per-fragment method, CHiCAGO. Combining inference across these two approaches is more stringent - a prey fragment needs to simultaneously have a higher count than expected and a supporting pattern among neighbouring fragments – and leads to improved resolution of direct contacts in CHi-C datasets.

## METHODS

### Pre-processing of read count data

We first pre-process the read count data using similar methods to standard CHi-C analysis to produce residuals which have a standard normal distribution in the absence of interactions. The raw data for a CHi-C experiment takes the form of a sparse matrix of counts for pairs of baits and preys. In practice, most entries in this matrix are zero, and analysis focuses on modelling the counts at preys that are within some linear genomic distance of each bait. Statistical inference of contacts is based on a two step approach. First, counts are modelled to adjust for systematic effects such as distance between bait and prey, and capture efficiency using either negative binomial (NB) regression (Dryden et al., 2014) or a convolution of NB and Poisson regression, to model biological and technical noise separately (Cairns et al., 2016). Second, a decision is taken to call contacts based on comparing observed counts to those expected under this empirical model estimated under a null hypothesis of no true interactions, either using raw p values (Dryden et al., 2014) or p values weighted to allow for the complication that we expect to find more interactions among fragments proximal to the bait, but test many more long distance pairs. We wished to use the first part of this procedure to account for the systematic effects in the data, and generate standardised residuals (that is, residuals with unit variance) for input into our proposed joint model.

The CHi-C data from CD4^+^ cells that we propose to use for this study have previously been processed by CHiCAGO (Cairns et al., 2016), and we noted that the technical noise component had a generally small contribution compared to the biological noise (Javierre et al., 2016) (Supplemental Fig S7). We therefore applied NB regression alone to the raw counts using standard software to generate these standardised residuals, which we call NB residuals in the text below. This allowed us to add additional covariates to the regression which we found provided small improvements to the model fit. Besides the distance between an interaction’s bait and prey fragments, we used both of their lengths, as well as transchromosomal bait activity, as covariates. Assuming that transchromosomal contacts are equally rare across baits (Johanson et al., 2017), the latter is a proxy for enrichment and capture biases. To account for the difference in the number of possible transchromosomal interaction sites between baits on different chromosomes, transchromosomal bait activity is defined for each bait as the residual following from the regression of the sum of its transchromosomal counts against its chromosome number. We used the R package GAMLSS is to fit zero-truncated NB models to counts for each pair of bait (*b*) and prey (*p*) within ten distance bins (Additional File 1, Table S3). Assuming most bait-prey fragment pairs do not make direct contacts, the null model can be parametrized using the full dataset. We used normalized, randomized quantile residuals (Dunn and Smyth, 1996) as the input for our joint model. A comparison between the predicted fits from CHiCAGO and our NB models applied to the same data showed a good correspondence (Supplemental Fig S8).

### Priors on model parameters

For each bait *b*, the non-zero strengths were assigned independent normal priors centered on 0, with a common variance 
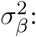

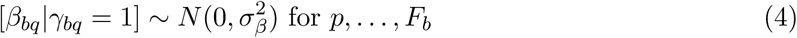

Rather than fixing *σ_β_*, which controls the magnitude of interaction strengths supported by the model, and therefore can have an important impact on the efficiency of the algorithm, we use a flexible hyper-prior allowing adaption to the data. Specifically, we placed a weakly informative Uniform(0.01,2) hyper-prior on *σ_β_*. The median,*σ_β_* = 1, corresponds to 95% support for interaction strengths up to a plausible 1.96. However, this hyper-prior equally supports much smaller values of *σ_β_*, as well as values up to the maximum of 2, corresponding to support for interaction strengths as large as 8 – marginally larger than any individual NB residual we observed (Supplemental Fig S1).

Our model selection framework is completed by specifying a prior for each *γ_bp_*. To avoid problems of over-fitting from simultaneous estimation of too many interaction parameters, and because we believe a *priori* that direct contacts only exist at a small proportion of prey fragments, our prior on *γ_bp_* is designed to encourage a “sparse” selection of interactions. To this end we first de ne *θ_b_*, the expected proportion of prey fragments which contact *b*, i.e. 
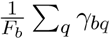
, which has prior distribution

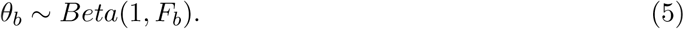

Conditional on *θ_b_*, each *γ_bq_* is then *i.i.d.* Bernoulli(*θ_b_*). This prior has two attractive properties. First, the marginal prior odds that a particular fragment interacts with *b* is 1/*F_b_*, and therefore decreases with the total number of fragments considered. Meanwhile, the prior odds for there being no interactions is a constant 0.5 for every bait. This setup provides an intrinsic multiplicity correction for the number of fragments in each bait, and allows fair comparison of inference across baits, due to the common prior on the null model (Wilson et al., 2010). Note too that this corresponds to a very small prior odds of interaction for each individual fragment, since *F_b_* is usually in the order of 3000, and thereby encourages the exploration of sparse models.

### Model Fitting via Reversible Jump MCMC

For bait *b*, the Reversible Jump MCMC (Green, 1995) sampling scheme starts at an initial set of interactions, *γ_b_*, and corresponding strengths, *β_b_*, denoted *γ_b_*(0) and *β_b_*(0) respectively. Bold symbols are used to denote that these are vectors across all fragments in the neighbourhood of bait *b*. To sample the next set of interactions and strengths, which we denote *γ_b_*(1) and *β_b_*(1), we propose moving from the current state to another combination of interactions and/or set of strengths, *γ_b_* * and 
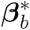
, by using a proposal function
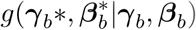
. We then accept these proposed values as the next sample with probability equal to the Reversible Jump Metropolis-Hastings ratio:

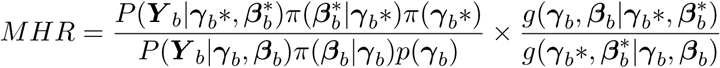

where ***Y**_b_* are the residuals of all fragments captured for bait *b*, *P*(***Y**_b_*|.) is the model described by (1) and (3). π( *β_b_*|*γ_b_*) is the prior distribution of the strengths defined in (4) conditional on the corresponding interactions being included in the model. *p*(*γ_b_*) is the model space prior dened in (5). Therefore the proposed combination of interactions and new strength values are accepted with a probability proportional to their likelihood and prior. If this new set of values is accepted, the proposed set is accepted as *γ_b_*(1) and *β_b_*(1); otherwise, the sample value remains equal to the current sample value, i.e., *γ_b_*(1) = *γ_b_*(0) and *β_b_*(1) = *β_b_*(0). It can be shown that this produces a sequence of parameter samples that converge to the required posterior distribution (Green, 1995). We used the Reversible Jump MCMC implementation in R2BGLiMS (https://github.com/pjnewcombe/R2BGLiMS, Newcombe et al. (2014)), to fit the model described above to each bait in turn. Because our aim was to fit this model to 28,214 baits, we took some time to define a strategy for thinning these samples in order to perform reliable posterior inference on *γ_b_* while minimising the computational burden (Additional File 2, Section 2). This led us to run R2BGLiMS sampling 5000 models per chain, at a density of 1 per 1000 iterations, with no burn-in. We ran two parallel chains for each bait, and checked convergence between MPPC derived from each chain. If this was < 0.75, we ran a further 5000 samples to improve convergence. Autocorrelation plots were also used to evaluate model space exploration for individual baits (Supplemental Fig S9).

### Assessing relationship of CHiCAGO scores and MPPC to outcome measures

CHiCAGO scores are non-negative real numbers, and are typically asinh transformed for presentation or downstream inference, to prevent over-leverage of points in the extreme right of the distribution (Javierre et al., 2016). In constrast, MPPC lies between 0 and 1, although rarely reaches 1 in practice. We found MPPC were generally positively correlated with CHiCAGO scores, with the relationship closest to linear when sqrt(MPPC) was compared to asinh(CHiCAGO) (Supplemental Fig S4). We therefore use a square root transform in following analyses to perform a fair comparison with CHiCAGO scores.

We defined the following four outcome measures:

**Promoter: match to Cao et al. (2017)** For validation with external promoter-enhancer networks, we used the positions given in http://yiplab.cse.cuhk.edu.hk/jeme/encoderoadmap_lasso/encoderoadmap_lasso.34.csv (accessed 2017/09/11). We used GenomicRanges to identifying bait-prey fragment pairs which overlapped the paired co-ordinates given in this file, and set a binary outcome 1 if such an overlap was found and 0 otherwise. Analyses of this measure were restricted to prey fragments within 200 kb of the bait, because 95% of these reported links were within that range.

**Promoter: link to active chromatin** These cells had previously been assayed by ChIP-seq, and a 15 state CHROMHMM model fitted. 8 of these states showed characteristics of “active chromatin” and we combined these into a binary measure for active or inactive chromatin (Burren et al., 2017b). We used these results to quantify the overlap, for each prey fragment, with regions of active chromatin. For the most part (∼ 90%), a fragment showed complete overlap or lack of overlap with active chromatin regions, in which case the outcome measures was set to 1 or 0 respectively. To allow logistic regression of this mainly binary outcome, the observations with fractional overlap were set to missing for analysis.

**Validation: overlap baited promoter** For a measure of promoter overlap, we used the binary indicator of whether a prey fragment in the validation experiment had also been baited in the promoter experiment.

**Validation: expression at linked promoter** Given evidence that recruitment of prey fragments associated with increased expression of the baited gene (Burren et al., 2017b), we expected that, amongst prey that did correspond to a baited promoter in the promoter capture experiment, the level of expression of the target gene should be higher when there was a direct contact. RNA-seq has previously been used to quantify transcription in these cells, and we used the expression of the target gene (log_2_(count + 1)) as an outcome measure in linear regression. Analyses of this measure were restricted to bait-prey pairs where the prey corresponded to a gene promoter.

Because each prey fragment is represented multiple times (with different baits), we assessed the relationship between asinh-transformed CHiCAGO scores and sqrt-transformed MPPC with each outcome measure using robust clustered linear or logistic regression implemented in the R library rms (https://cran.r-project.org/web/packages/rms/), clustering on the prey fragment.

## DATA ACCESS

Data used is available as described in the primary publications. CHi-C data, inferred CHROMHMM states, RNA-seq quantification are available as described in (Burren et al., 2017b). We downloaded summaries of the enhancer-promoter networks (Cao et al., 2017) http://yiplab.cse.cuhk.edu.hk/jeme/encoderoadmap_lasso/encoderoadmap_lasso.34.csv.

The processed CHi-C data, with CHiCAGO scores, NB residuals and MPPC statistics are in supplementary data supp-data.tgz. This gzipped archive contains four .csv les (suppdata-*.csv), one for each experiment, with bait and prey *hind*III fragment IDs together with columns:

**N** read count

**residual** NB residual

**mppc** MPPC

**beta.post** posterior expectation of *β*

**chicago CHiCAGO** score and two annotation files: `hind-positions.csv` gives the chromosome co-ordinates of each *hind*III fragment and `bait2gene.csv` gives the links between promoter baits and annotated genes, using ensembl 75.

**NOTE** We failed to upload the supporting supplementary data several times to the journal. Instead, please find it at https://www.dropbox.com/s/wny9m1gtcxefrls/supp-data.tgz?dl=0 We provide functions to implement this model in the R package Peaky, available from http://github.com/cqgd/pky.

Code used to generate the tables and figures in this paper are available at https://github.com/chr1swallace/eijsbouts-et-al

## ACKNOWLEDGEMENTS

We thank Frank Dudbridge and Mikhail Spivakov for helpful discussions throughout the development of our method.

This work was funded by the MRC (MC UP 1302/5, MC UP 0801/1) and the Wellcome Trust (WT107881).

## Author’s contributions

CE developed the statistical method, implemented it, applied it to the CHi-C datasets, wrote the paper, authored the software. OB annotated gene TSS and CHi-C *Hind*III fragments, critically evaluated the enrichment analysis, wrote the paper. PN developed the statistical method, wrote the paper. CW devised the study, developed the statistical method, performed statistical enrichment analyses, wrote the paper.

## DISCLOSURE DECLARATION

The authors declare that they have no competing interests.

